# Comparison of Dimensionality Reduction and Clustering Methods for Single-Cell Transcriptomics Data

**DOI:** 10.1101/2022.10.15.512334

**Authors:** Vrushali Pandit, Asish Kumar Swain, Pankaj Yadav

## Abstract

Dimensionality reduction (DR) methods are applied to extract relevant features from inherently high dimensional and noisy single-cell RNA sequencing (scRNA-seq) data. Choice of DR method could influence the performance of clustering algorithm and subsequent analysis outcomes. We performed a benchmarking study of seven popular DR methods and four clustering algorithms widely used for scRNA-seq datasets. For this purpose, we used three publicly available real scRNA-seq datasets. The performance was evaluated using two clustering metrics viz. adjusted random index (ARI) and normalized mutual index (NMI). We also compared our results with a similar study published by Xiang and colleagues. Overall, we observed higher ARI and NMI scores for DR methods when compared with Xiang’s study. We also noticed several differences between our and Xiang’s study. Noteworthy, three methods, namely, Independent Component Analysis (ICA), t-Distributed Stochastic Neighbor Embedding (t-SNE) and Uniform Manifold Approximation and Projection (UMAP) performed consistently well across three datasets. Linear method ICA was best performer on *Segerstolpe* dataset, while nonlinear methods UMAP and t-SNE best performed on *Deng* and *Chu* datasets, respectively. Neural network-based methods Variational Autoencoder (VAE) and Deep Count Autoencoder (DCA) could not perform well probably due to their sensitivity to hyperparameters and overfitting. Among clustering methods, Gaussian Mixture Models (GMMs) performed consistently well across datasets. This might be because GMMs are the universal approximators of posterior probability densities. We conclude that performance of different DR methods is more dataset dependent and for various scRNA-seq datasets different algorithms are more suited and there is no one-fit-all method.

## Introduction

Conventionally, cell-based research has relied on bulk RNA sequencing for various purposes such as interpretation of the functional elements of the genome, identification of molecular constituents of cells, and understand development and diseases. One major limitation of the bulk RNA sequencing approach is that it captures only the average transcriptomic signal of cells collected from a tissue sample. Recent advancements in the field of fluorescence-activated cell sorting, magnetic activated cell sorting, laser capture microdissection, microfluidics, and sequencing technologies have made it plausible to capture individual cells including rare cell types [1]–[5]. Now, it is possible to obtain the transcriptome data for individual cells [6]. In the past few decades, the amount of single cell RNA sequencing (scRNA-seq) data in public databases has increased exponentially. In the year 2017, the *10X Genomics*company released a dataset of 1.3 million cells publicly available to researchers [7]. In theory, the scRNA-seq technologies can generate whole transcriptome data at single cell resolution which is intrinsically difficult to analyze due to its high dimensionality. Typically, the scRNA-seq data consists of approximately 20,000 features (genes) and up to one million observations (cells). In addition, the cellular subpopulations consisting of various tissues exhibit continuous gene expression changes related to cell cycle progression, spatial coordinates, and differences in differentiation maturity. This heterogeneity further increases the dimensionality of scRNA-seq data.

Dimensionality reduction methods are used to extract meaningful features which could be utilized further for downstream analyses such as clustering, trajectory analysis, rare cell type annotation, and data integration [8]. For instance, to identify cell types in a tissue sample, the clustering of similar cells is performed in an unsupervised manner using the latent space generated by a dimensionality reduction method. The clustering of cells requires the ability to detect relationships and maximize internal density and external sparsity. Various clustering methods are available, but only a handful of them are useful for the complicated and highly non-linear scRNA data. The k-means clustering is the most widely used algorithm, but it can only cluster linearly separable data [9]. There are other types of clustering methods based on the probabilistic distribution. Of these, hierarchical agglomerative clustering is a bottom-up approach that keeps aggregating small and similar clusters into a tree. The gaussian mixture model (GMM) provides the best approximation of the posterior distribution[10]. Spectral clustering relies on graph theory, which uses a similarity matrix of the data to perform dimensionality reduction before performing clustering at lower dimensions [11], [12]. The performance of clustering methods relies on the outcome of the dimensionality reduction method. In literature, several dimensionality reduction methods are available which have been categorized as linear, nonlinear, model-based, and neural network based. The most widely used linear dimensionality reduction method is principal component analysis (PCA) [13]. This is an unsupervised method that retains only those linear combinations of original features which maximize variance along the entire data. PCA only retains combinations of features that maximize variance. The independent component analysis (ICA) is another linear dimensionality reduction method. The ICA method minimizes the dependencies among new features instead of maximizing the variance. Two popular non-linear models for dimensionality reduction include t-distributed student neighbor embedding (t-SNE) and uniform manifold approximation and projection (UMAP) [14], [15]. The zero-inflated factor analysis (ZIFA) is a model-based dimensionality reduction method that explicitly accounts for the occurrence of dropout events which usually occur in scRNA-seq datasets [16]. Among neural network-based dimensionality reduction methods, the variational autoencoders (VAE) and deep count autoencoders (DCA) have recently gained attraction for scRNA-seq datasets [17], [18].

The utility of different dimensionality reduction methods and clustering algorithms for scRNA-seq data has not been fully explored so far. Recently, two benchmarking studies have been reported in the literature focusing on the dimensionality reduction methods [19], [20]. In one study, Xiang and colleagues compared the performance of 10 dimensionality reduction algorithms using 10 simulated and 5 real scRNA-seq data. However, their study considered only k-mean clustering algorithm and did not cover other useful approaches for scRNA-seq data. Since k-means might not be efficient for complicated scRNA-seq data, it might be worthwhile exploring the utility of other clustering approaches as well. Another benchmarking study by Tsuyuzaki and colleagues was limited only towards the performance of faster and memory efficient PCA algorithms. Again, PCA methods are simplistic and linear model and might not be robust for scRNA-seq data. Thus, there is an immediate need for a comprehensive benchmarking of important dimensionality reduction methods as well as clustering algorithms available for the scRNA-seq data.

In this study, we benchmark seven different dimensionality reduction methods, namely, PCA, ICA, t-SNE, UMAP, ZIFA, VAE and DCA using three distinct real scRNA-seq datasets. Moreover, we benchmarked four relevant and distinct clustering algorithms, namely, k-means, agglomerative, GMM and spectral clustering to assess their ability to identify similar cell types on reduced feature space obtained from dimensionality reduction methods. The performance of these methods is evaluated based on how well they could extract cell-to-cell variability measured based on two simple metrics.

## Materials and Methods

### scRNA-seq Datasets

We used three publicly available scRNA-seq datasets to benchmark the performance of dimensionality reduction methods and clustering algorithms. The same datasets were also used by Xiang’s paper and are summarized in **Table 1.** These datasets were obtained from different species and comprise diverse cell types. Of these, the *Segerstolpe* dataset contains the RNA profiling of singular cells of the human pancreatic tissue obtained from healthy individuals and type 2 diabetic patients [21]. This dataset is most diverse and contains 14 cell types. It also has the highest number of 2209 cells and 26,271 genes compared to the other two datasets. The *Deng* dataset contains RNA profiling across individual cells of mouse-preimplantation embryos of mixed backgrounds [22]. This dataset comprised a total of 269 cells of 6 cell types, and 22,958 genes. The *Chu* dataset contains transcriptomic data of individual pluripotent stem cells [23]. This dataset comprised 1018 cells of 7 cell types and 19,097 genes.

**Table 1.**
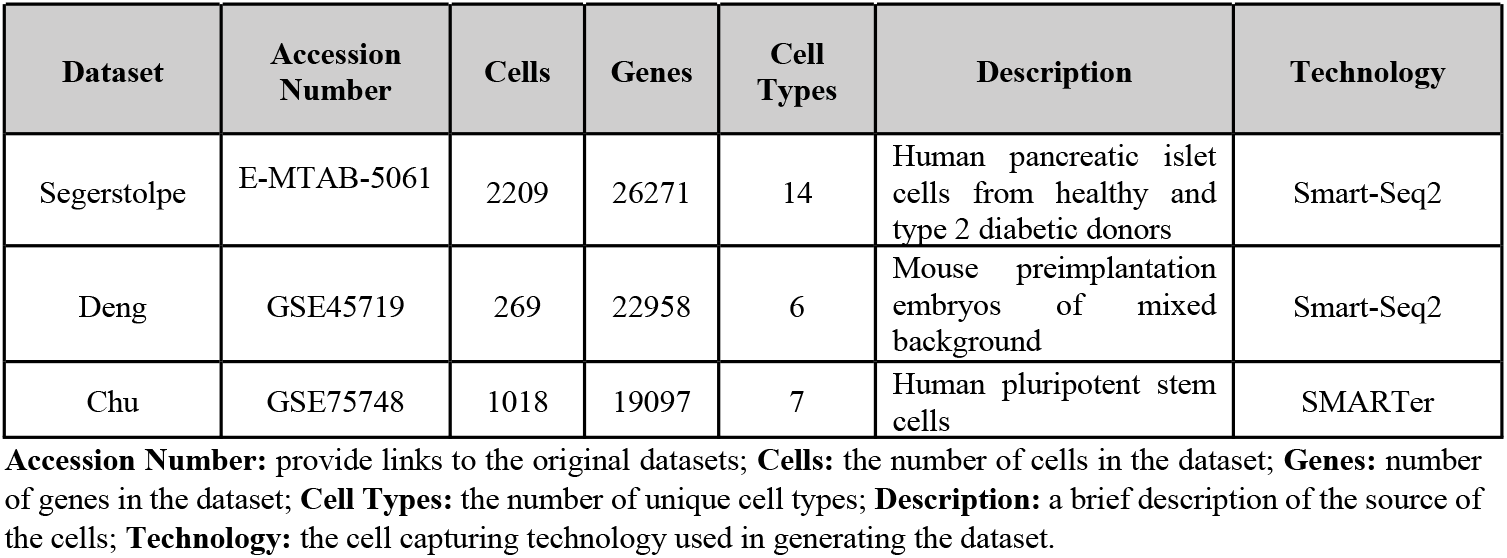
Summary of three scRNA-seq datasets used in this benchmarking study.

| Dataset | Accession Number | Cells | Genes | Cell Types | Description | Technology |
| --- | --- | --- | --- | --- | --- | --- |
| Segerstolpe | E-MTAB-5061 | 2209 | 26271 | 14 | Human pancreatic islet cells from healthy and type 2 diabetic donors | Smart-Seq2 |
| Deng | GSE45719 | 269 | 22958 | 6 | Mouse preimplantation embryos of mixed background | Smart-Seq2 |
| Chu | GSE75748 | 1018 | 19097 | 7 | Human pluripotent stem cells | SMARTer |

**Accession Number:** provide links to the original datasets; **Cells:** the number of cells in the dataset; **Genes:** number of genes in the dataset; **Cell Types:** the number of unique cell types; **Description:** a brief description of the source of the cells; **Technology:** the cell capturing technology used in generating the dataset.

### Data Analysis

Our analysis workflow is illustrated in the **Figure 1**. The three scRNA-seq datasets were available to us in different formats. At first, we converted all these datasets into a matrix which can be utilized further by various dimensionality reduction methods. For the *Segerstolpe* dataset, we removed the empty droplets as well as droplets containing doublets (i.e., 2 cells). Next, we mapped each cell to corresponding cell types using unique identifiers which were provided with individual datasets. For the *Deng* dataset, we combined individual cell data stored in different text files into a single matrix. In some datasets both raw count data as well as RPKM (reads per kilobase data was available of transcript per million) were available. As RPKM is normalized for both sequencing depth and the gene length, we used RPKM instead of raw count data whenever it was available. The *Segerstolpe* and *Deng* datasets were normalized using standard scaling. The Chu dataset was already normalized using a median-by-ratio normalization, hence there was no need for further normalization. The normalized datasets were used as input for different dimensionality reduction methods.

**Figure 1.**
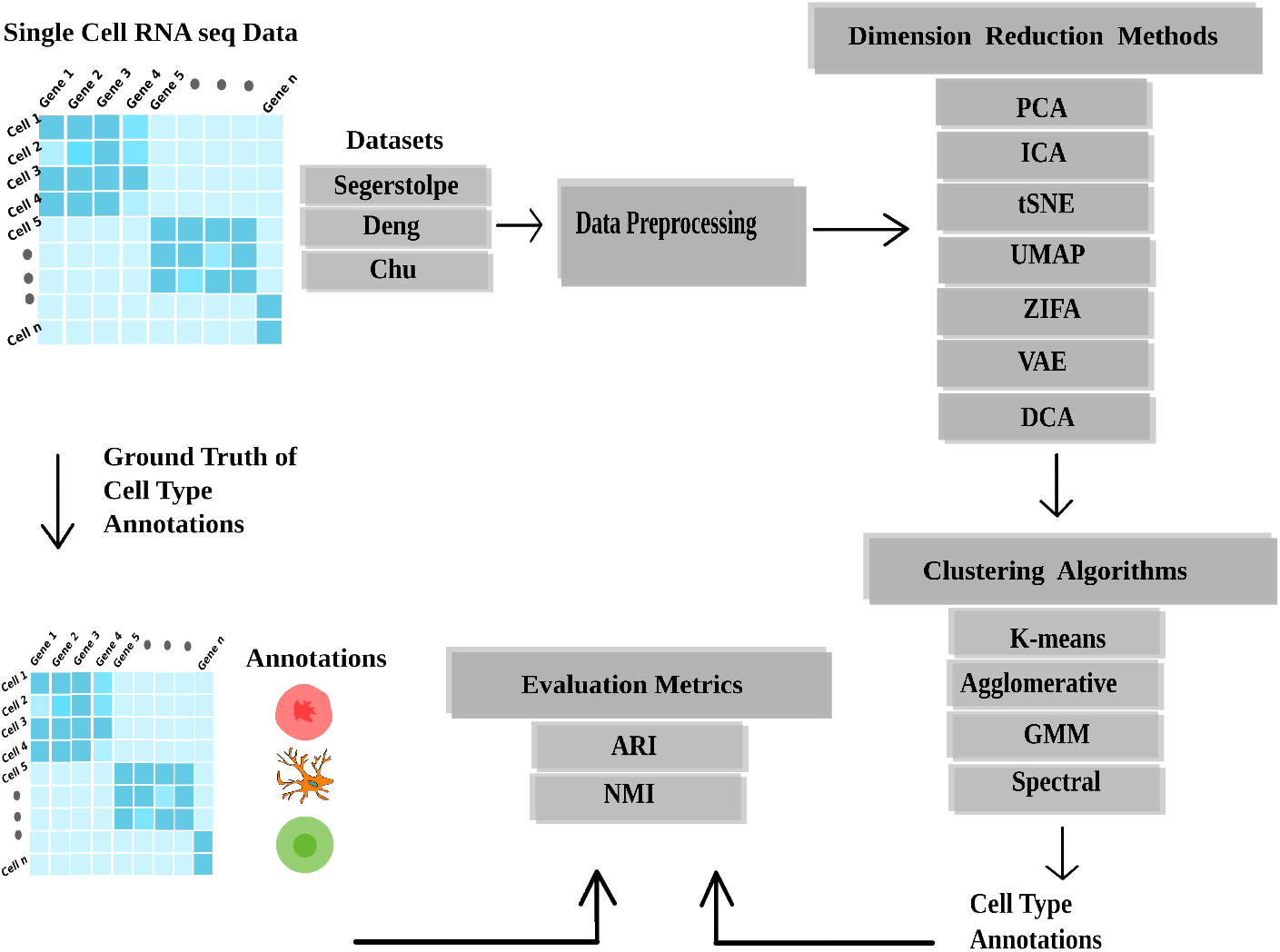
Show the analysis workflow followed in this benchmarking study. Three real scRNA-seq datasets were used and the actual cell types were known. The read count values were normalized either using reads per kilobase of transcript per million reads mapped (RPKM) or median-by-ratio method. After data preprocessing and normalization, dimensionality reduction was performed using 7 different methods for each dataset. Clustering analysis was performed in an unsupervised manner using 4 different methods. The performance of dimensionality reduction algorithms was assessed based on the k-means clustering accuracies. The performance of clustering algorithms was assessed based on accuracies of clusters obtained from best performing dimensionality reduction algorithm for each dataset

### Dimensionality Reduction Methods

We benchmarked the performance of seven different dimensionality reduction methods which included 2 linear, 2 non-linear, one model-based and 2 neural network-based methods (see **Table 2**). These methods are either generic or specific to the scRNA-seq data and briefly described below.

**Table 2.** Summary of seven different dimensionality reduction algorithms benchmarked in this study.

| Algorithm | Category | Advantages | Disadvantages |
| --- | --- | --- | --- |
| Principal component analysis (PCA) | Linear | Removes all redundant highly correlated features as all the principal components are independent of each other | The main disadvantages are data leakage and loss of interpretability of new PCs |
| Independent component analysis (ICA) | Linear | The main advantage of using ICA is that it is able to decompose mixed signals into independent sources. | The main disadvantage lies in its assumptions. |
| t-student stochastic neighbor embedding (t-SNE) | Nonlinear | The main advantage of using t-SNE over other methods is that it can handle nonlinear data which has complex local feature relationships very well. | The bottleneck with using this method is that it is computationally expensive, time consuming and cannot preserve global features. |
| Uniform Manifold Approximation and Projection (UMAP) | Nonlinear | Addresses all the shortcoming of tSNE. tSNE is more of a visualization tool, but UMAP can be used as a general-purpose dimensionality reduction tool. | The only disadvantage with UMAP is that as it is a fairly new algorithm, it is not optimized. |
| Zero Inflated Factor Analysis (ZIFA) | Model based | One of the only DR algorithms which explicitly accounts for the occurrence of dropout events. | Some of the main disadvantages of ZIFA are that it takes very long to execute (which has limited its application to large single cell datasets) and that it only works with non-normalized raw count data. |
| Variational Autoencoder (VAE) | Neural network | VAE has more parameters to tune that gives significant control over how we want to model our latent distribution | A disadvantage of VAEs is that the generated samples are blurred due to injected noise and imperfect reconstruction. |
| Deep Count autoencoder (DCA) | Neural network | DCA allows the user to specify the noise model. This gives the model high scalability. | The fixed neuronal architecture of DCA can be restrictive. |

#### 1. Principle Component Analysis (PCA)

This is a linear method which uses independent linear combinations of original feature vectors and arranges them in descending order of their variance. Then depending on either the number of components decided by the user or percentage of variance captured, these top components become the new feature space. The main advantage of using PCA is that it removes redundant highly correlated features, and all the principal components are independent of each other. The main disadvantages of PCA are data leakage and loss of interpretability of new PCs [13].

#### 2. Independent Component Analysis (ICA)

Like PCA this is also a linear method. However, instead of maximizing the variance, ICA minimizes dependencies among new features. Two assumptions of ICA are that the source signals should be independent of each other and should not follow a gaussian distribution. PCA is a data compressor, while ICA is more of a data separator. The main advantage of using ICA is that it can decompose mixed signals into independent sources. The main disadvantage lies in its assumptions [24].

#### 3. t-Stochastic Neighbour Embedding (t-SNE)

t-SNE is an unsupervised nonlinear method for data visualization. It calculates a similarity measure between pairs of instances in the high dimensional space based on a gaussian kernel and in the low dimensional space based on a heavy tailed t-student kernel distribution. It then tries to optimize these two similarity measures using a cost function. The main advantage of using t-SNE over other methods is that it can very well handle nonlinear data with complex local feature relationships. The bottleneck with using this method is that it is computationally expensive and cannot preserve global features [14].

#### 4. Uniform Manifold Approximation and Projection (UMAP)

Like t-SNE, this method also uses graph layout algorithms to arrange data in low-dimensional space. This method is faster compared to t-SNE. Moreover, it can better preserve global relationships and can deal with large data. UMAP constructs a high dimensional graph representation of the data then optimizes a low-dimensional graph to be as structurally similar as possible. The underlying mathematics behind this is based on fuzzy simplicial complex [15].

#### 5. Zero Inflated Factor Analysis (ZIFA)

Due to the high level of dropouts and sparsity in the scRNA-seq data, traditional methods may seem to not work well. This method explicitly models the dropout characteristics. ZIFA adopts a latent variable model based on the factor analysis framework and augments it with an additional zero-inflation modulation layer. The main disadvantages of ZIFA are that it takes very long to execute (which has limited its application to large single cell datasets) and that it only works with non-normalized raw count data [16].

#### 6. Variational Autoencoder (VAE)

This is a modification of the traditional autoencoder. However, instead of mapping the data to a fixed bottleneck layer, VAE maps it to a distribution with a mean and a standard deviation layer. Sampling with a reparameterization trick is done from these two layers to allow for backpropagation to occur. A disadvantage of VAEs is that the generated samples are blurred due to injected noise and imperfect reconstruction [17].

#### 7. Deep Count Autoencoder (DCA)

This is a special architecture of the vanilla autoencoder. The deep learning framework (by default three hidden layers with 64, 32, 64 neurons) of DCA enables the capturing of the complexity and non-linearity in scRNA-seq data. To provide maximal flexibility, DCA implements a selection of scRNA-seq specific noise models including negative binomial distribution with (ZINB) and without zero-inflation (NB). For example, using the ZINB noise model, DCA learns gene-specific parameters mean, dispersion and dropout probability based on the input gene expression data [18].

### Clustering Algorithms

Usually in experiments, the cell types of individual cell samples are unknown a priori, and hence a robust clustering algorithm is used to identify cell types. The performance of clustering method relies majorly on the outcome of the dimensionality reduction method. For each dataset, we performed k-means clustering on the reduced feature space obtained from the dimensionality reduction methods. The performance of dimensionality reduction method was assessed based on how well the reduced data could retain information regarding the cell types.

Moreover, we evaluated four clustering algorithms with different underlying logic such as centroid based (k-means), graph partitioning based (spectral clustering), posterior distribution based (gaussian mixture models) and hierarchical clustering (agglomerative). **Table 3** provides an overview of the four clustering methods evaluated in our work and are described below.

**Table 3.**
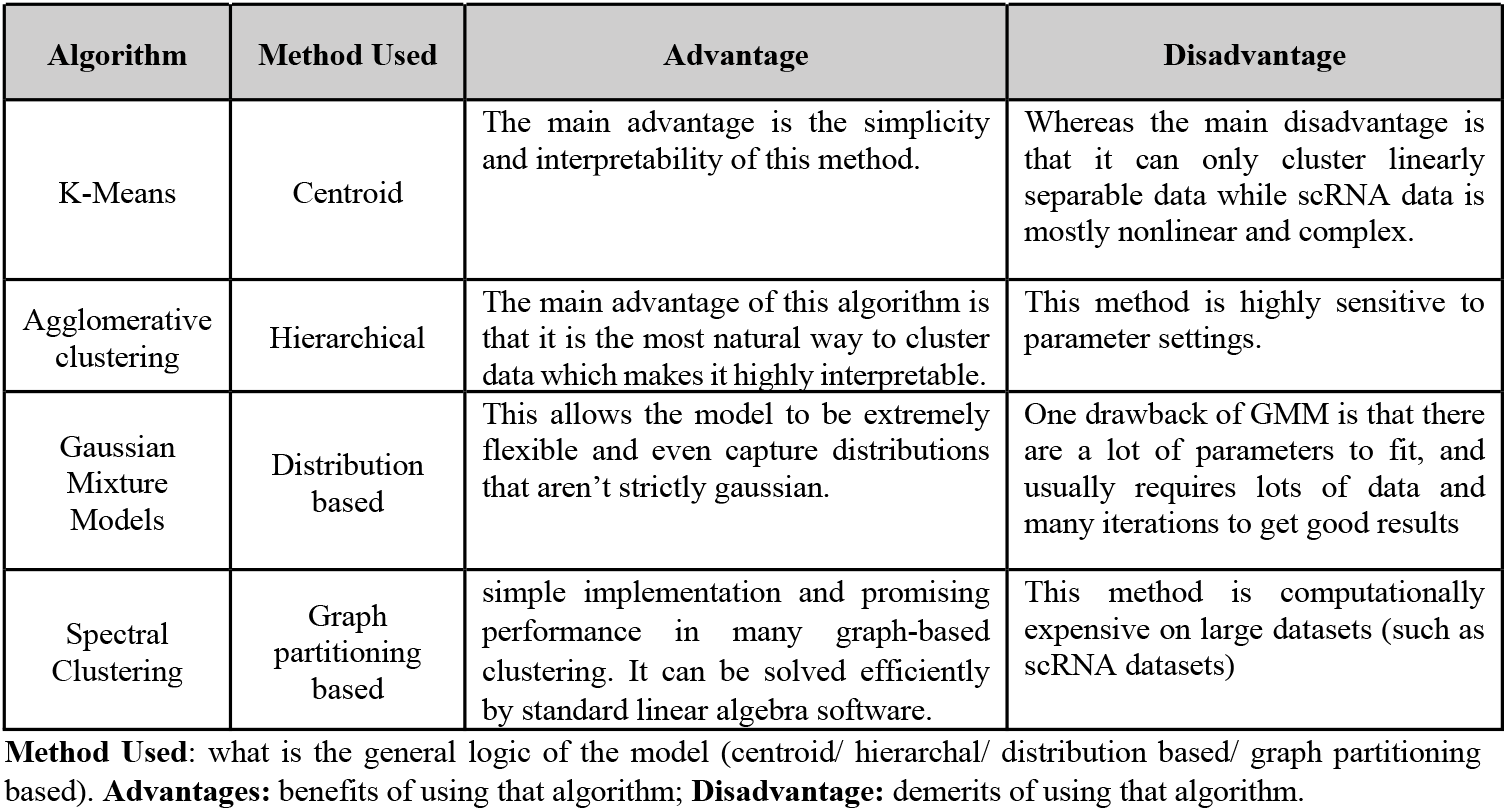
Summary of four clustering algorithms benchmarked in this study.

| Algorithm | Method Used | Advantage | Disadvantage |
| --- | --- | --- | --- |
| K-Means | Centroid | The main advantage is the simplicity and interpretability of this method. | Whereas the main disadvantage is that it can only cluster linearly separable data while scRNA data is mostly nonlinear and complex. |
| Agglomerative clustering | Hierarchical | The main advantage of this algorithm is that it is the most natural way to cluster data which makes it highly interpretable. | This method is highly sensitive to parameter settings. |
| Gaussian Mixture Models | Distribution based | This allows the model to be extremely flexible and even capture distributions that aren't strictly gaussian. | One drawback of GMM is that there are a lot of parameters to fit, and usually requires lots of data and many iterations to get good results |
| Spectral Clustering | Graph partitioning based | simple implementation and promising performance in many graph-based clustering. It can be solved efficiently by standard linear algebra software. | This method is computationally expensive on large datasets (such as scRNA datasets) |

**Method Used:** what is the general logic of the model (centroid/ hierarchal/ distribution based/ graph partitioning based). **Advantages:** benefits of using that algorithm; **Disadvantage:** demerits of using that algorithm.

#### 1. K-means Clustering

K-means clustering is one of the simplest and most intuitive clustering algorithms. Initially, the random datapoints are assigned as cluster centers. All the samples are assigned to a cluster center based on minimizing a similarity function of choice such as Euclidean distance. The centroids of so formed clusters become the new cluster centers. This continues till convergence. The main advantage of this method is the simplicity and interpretability, while the main disadvantage is that it can only cluster linearly separable data while scRNA data is mostly nonlinear and complex [25].

#### 2. Agglomerative Clustering

This method follows a bottom-up hierarchical approach. Initially, every datapoint is treated as a cluster. Later based on various types of linkages (such as average, complete, single) and similarity measures (such as Euclidean, Manhattan, cosine distance) all the clusters are grouped together up into larger clusters to a threshold set by the user. The main advantage of this algorithm is that it is the most natural way to cluster data which makes it highly interpretable. However, this method is highly sensitive to parameter settings [26].

#### 3. Gaussian Mixture Model (GMM)

This method is also known as the universal approximator of distributions which is analogous to the role of neural networks being the universal approximators of functions. It does probabilistic soft assignment of clusters which often leads to much better results than naive hard assignment. Three parameters must be learned includes mean, covariance and a mixing probability that defines how big or small the gaussian function will be. These parameters are found using the EM algorithm [10].

#### 4. Spectral Clustering

This algorithm uses graph theory, where communities of nodes are identified in a graph based on the edges connecting them. It first creates a similarity graph between all the data points to cluster. This graph can be made by representing various relationships between data points like – an epsilon neighborhood graph or a kNN graph. It then computes the eigenvectors of the graph’s Laplacian matrix. The first k eigen vectors are then clustered using KMeans. The method is flexible and allows us to cluster non-graph data as well [11].

### Evaluation Metrics

The performance of clustering algorithm was assessed using two metrics, namely, adjusted random index (ARI) and normalized mutual index (NMI) [27], [28]. The dimensionality reduction method with highest average of ARI and NMI scores was considered as the best performer for the given dataset. These both metrics provide a measure of agreement between two partitions by calculating similarity between true cell type labelling and clustering labelling. The ARI creates a contingency table from the clustered data which stores the co-occurrences of a sample in two different clustering including the ground truth and the result of dimensionality reduction and clustering algorithm. The ARI score ranges between 0 to 1 where a larger score means that two clusters are more consistent with each other. Conversely, the score 0 corresponds to randomly generated clusters. X and Y are defined as the community assignments for each node in the graph. Each pair of nodes of the graph i, j can be fit to one of four categories:

N_00_= i and j are assigned to different clusters in both X and Y

N_01_= i and j are assigned to different clusters in X but to the same cluster in Y

N_10_= i and j are assigned to the same cluster in X but to different clusters in Y

N_11_= i and j are assigned to the same cluster in both X and Y

The ARI is calculated using formula,

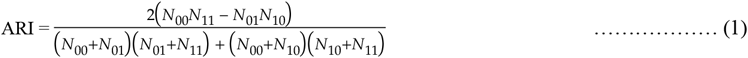

The NMI is a variant of a common measure in information theory called mutual information to scale the results between 0 (no mutual information) and 1 (perfect correlation). It is built on the Shannon entropy of information theory. Let partitions X and Y define community assignments {x_i_} and {y_i_} for each node *i* in the graph.

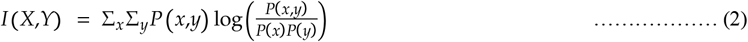

To normalize the I(X,Y) values, the following equation is used:

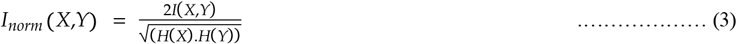

Where, H(X) = -∑x P(x)log P(x) is the Shannon’s entropy and P(x) is the probability that node picked at random is assigned to the community x.

## Results

### Benchmarking of Dimensionality Reduction Methods

We benchmarked the performance of 7 different dimensionality reduction methods, namely PCA, ICA, t-SNE, UMAP, ZIFA, VAE, and DCA using three diverse scRNA datasets. All three datasets contained standard scaled RPKM values of genes expressed by individual cells. For each dataset, we performed k-means clustering on the reduced feature space generated by the respective seven dimensionality reduction methods. The performance of each dimensionality reduction method was assessed based on how well the reduced data could retain information regarding the cell types. The clustering performance was evaluated using ARI and NMI metrics. The dimensionality reduction method with the highest average of ARI and NMI scores was deemed best performer for the given dataset. Moreover, the clustering accuracies obtained in our study were compared with their respective values reported by Xiang and colleagues [19] The results obtained for each dataset are described below.

#### 1. Segerstolpe dataset

Overall, we observed higher ARI and NMI scores as compared to those previously reported in Xiang’s paper for all dimensionality reduction methods except ZIFA (**Table 4**). For ZIFA method, the average ARI and NMI score was very close to the reported score in Xiang’s paper (**Figure 2A**). Noteworthy, the ICA was best performer among seven methods in our study with ARI and NMI scores of 0.88 and 0.84, respectively. To our surprise, the corresponding ARI and NMI scores for ICA method previously reported by the Xiang’s paper were much smaller (0.03 and 0.11, respectively). Moreover, Xiang’s paper reported VAE as the best performer with average ARI and NMI score 0.41 while our study observed an average score 0.49 for this method. **Figure 2B** shows the percentage increment in ARI and NMI scores for seven methods with respect to Xiang’s paper. We observed average ARI and NMI scores for PCA, t-SNE, UMAP, VAE and DCA increased by 2, 1.5, 2.3, 1.2, and 1.6. fold, respectively, relative to Xiang’s paper. We would like to mention that the *Segerstolpe* dataset originally has only RPKM data, however DCA requires raw count data. Hence, we re-normalized each feature by multiplying it with its gene length then multiplying them by the per million scaling factor.

**Figure 2.**
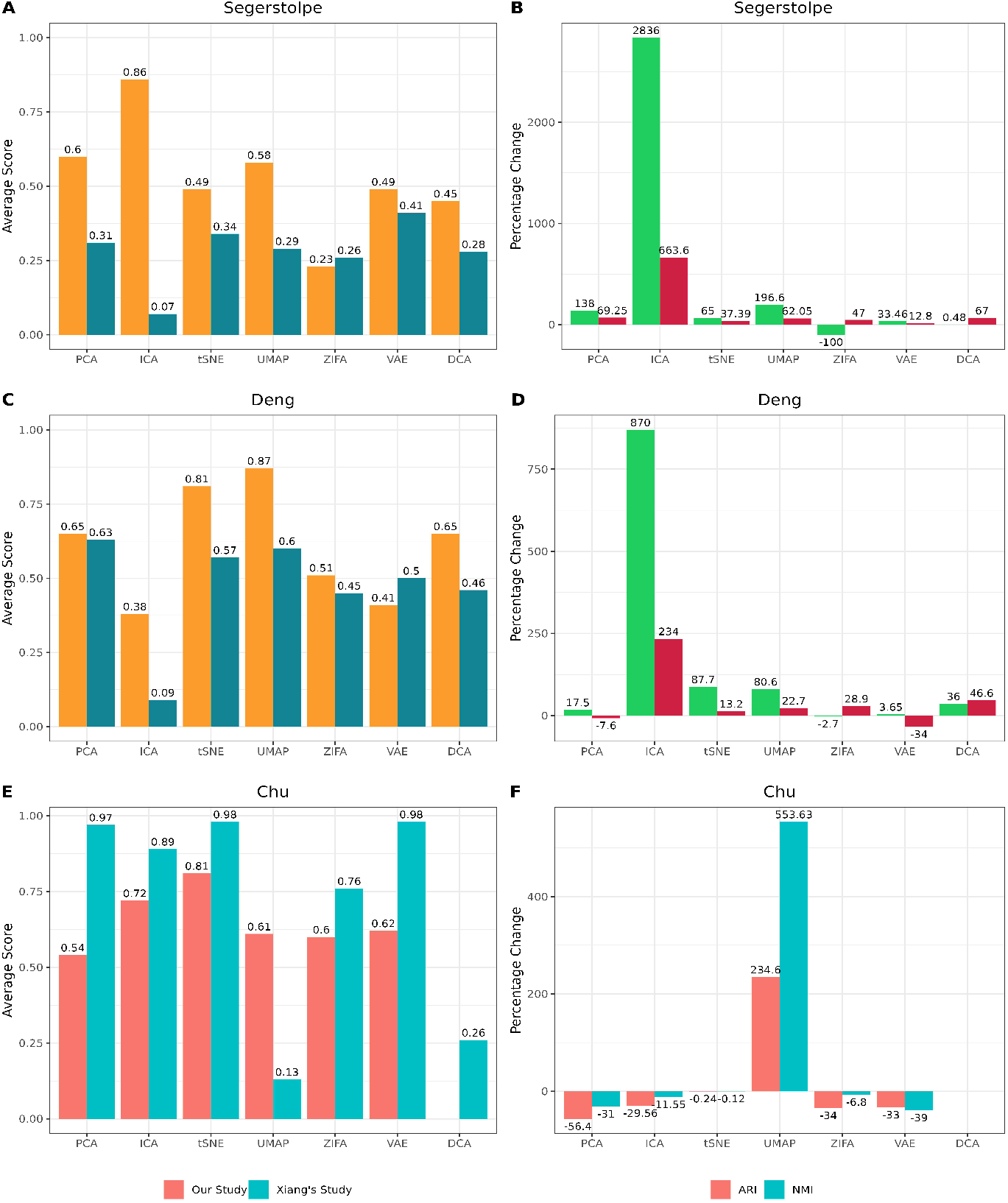
Histogram showing average ARI and NMI scores for seven dimensionality reduction methods considered in this study: panels A, C and E refers to Segerstolpe, Deng and Chu datasets, respectively. Also, shown is the percentage changes in the ARI and NMI scores relative to the corresponding values reported in Xiang’s paper. The panels B, D and E refers to Segerstolpe, Deng and Chu datasets, respectively.

**Table 4.** Comparison of the performance of seven dimensionality reduction (DR) methods based on ARI and NMI scores for three different datasets.

| DR Method | Segerstolpe Dataset |  |  |  | Deng Dataset |  |  |  | Chu Dataset |  |  |  |
| --- | --- | --- | --- | --- | --- | --- | --- | --- | --- | --- | --- | --- |
|  | ARI Score |  | NMI Score |  | ARI Score |  | NMI Score |  | ARI Score |  | NMI Score |  |
|  | Our Study | Published | Our Study | Published | Our Study | Published | Our Study | Published | Our Study | Published | Our Study | Published |
| PCA | 0.52 | 0.22 | 0.68 | 0.40 | 0.63 | 0.53 | 0.67 | 0.73 | 0.43 | 0.99 | 0.65 | 0.95 |
| ICA | 0.88 | 0.03 | 0.84 | 0.11 | 0.29 | 0.03 | 0.47 | 0.14 | 0.64 | 0.91 | 0.81 | 0.87 |
| t-SNE | 0.35 | 0.21 | 0.63 | 0.46 | 0.85 | 0.45 | 0.77 | 0.68 | 0.76 | 0.99 | 0.86 | 0.97 |
| UMAP | 0.53 | 0.18 | 0.63 | 0.39 | 0.89 | 0.49 | 0.86 | 0.70 | 0.50 | 0.15 | 0.72 | 0.11 |
| ZIFA | 0.00 | 0.19 | 0.47 | 0.32 | 0.42 | 0.43 | 0.61 | 0.47 | 0.51 | 0.78 | 0.69 | 0.74 |
| VAE | 0.35 | 0.26 | 0.63 | 0.56 | 0.43 | 0.41 | 0.39 | 0.59 | 0.65 | 0.99 | 0.59 | 0.97 |
| DCA | 0.34 | 0.23 | 0.55 | 0.33 | 0.56 | 0.41 | 0.75 | 0.51 | NA | 0.21 | NA | 0.31 |

**DR Method:** refers to dimensionality reduction method used in the study, **ARI Score:** refers to Adjusted Random Index score which is a clustering accuracy metric, **NMI Score:** refers to Normalized Mutual Index score which is another clustering accuracy metric; **Our study:** indicates the ARI and NMI scores obtained in this benchmarking study; **Published:** indicates the ARI and NMI scores previously reported in the Xiang’s paper.

#### 2. Deng dataset

Like *Segerstolpe* dataset, we found higher ARI and NMI scores in comparison to Xiang’s paper for all dimensionality reduction methods benchmarked in our study except for VAE method which produced slight lower scores (**Table 4** and **Figure 2C**). Here, UMAP performed the best with an average ARI and NMI score 0.87 followed by t-SNE with average ARI and NMI score 0.81. In contrast, Xiang’s paper reported PCA as the best performer with average ARI and NMI score 0.63. Like Segerstolpe dataset, we observed approximately seven-fold increase in the accuracy for ICA method with average ARI and NMI scores 0.65 and 0.09 observed in our study and reported Xiang’s study, respectively. **Figure 2D** shows the percentage change in ARI and NMI scores for each method with respect to Xiang’s paper.

#### 3. Chu dataset

For the *Chu* dataset, t-SNE method performed the best with an average ARI and NMI score 0.81 (**Figure 2E**). Interestingly, Xiang’s paper also reported t-SNE as one of the best performers in their study for this dataset with average ARI and NMI score 0.98. Here, in contrast to *Segerstolpe* and *Deng* datasets, we observed an opposite trend in the performance of five dimensionality reduction methods with respect to Xiang’s paper (see **Table 4** and **Figure 2E**). We observed average ARI and NMI scores for PCA, ICA, t-SNE, ZIFA and VAE decreased by 0.56, 0.82, 0.82, 0.80 and 0.63 fold, respectively, relative to Xiang’s paper (**Figure 2F**). The UMAP method showed the highest percentage increase in the ARI and NMI scores while PCA method showed the highest percentage decrease in the ARI and NMI scores. The DCA method could not be applied to this dataset as the original data available to us was median-by-ratio normalized while DCA requires raw counts.

### Benchmarking of Clustering Algorithms

We benchmarked the performance of 4 different clustering algorithms namely, k-means, agglomerative, GMM and spectral clustering. For each clustering algorithm, we utilized the feature space reduced by the best performing dimensionality reduction methods chosen based on ARI and NMI scores for each dataset. Thus, the ICA method extracted features were used for the *Segerstolpe* dataset. Similarly, the UMAP and t-SNE extracted features were used for the *Deng* and *Chu* datasets, respectively. We assessed the accuracy of these clustering algorithms based on how well they could identify cell types which were evaluated using the ARI and NMI metrics. We found considerable differences in the performance of the 4 clustering methods for the *Sergerstolpe* and *Deng* datasets, but for the *Chu* dataset the differences were only marginal (**Figure 3**). For the *Segerstolpe* dataset, the GMM method performed the best with an average score of 0.87 (**Figure 3A**). This was followed by k-means, agglomerative clustering, and spectral clustering with average scores 0.86, 0.51, and 0.34, respectively. Likewise, for the *Deng* dataset, the GMM method again performed the best with an average score of 0.76 followed by agglomerative clustering, spectral clustering, and k-means with average scores 0.54, 0.51, and 0.34 (**Figure 3B**). For the *Chu* dataset, again GMM performed the best with an average score of 0.84 followed by agglomerative clustering, spectral clustering, and k-means with average scores 0.82, 0.81, and 0.81, respectively (**Figure 3C**). Noteworthy, the GMM method was the best performer across all the three considered scRNA-seq datasets.

**Figure 3.**
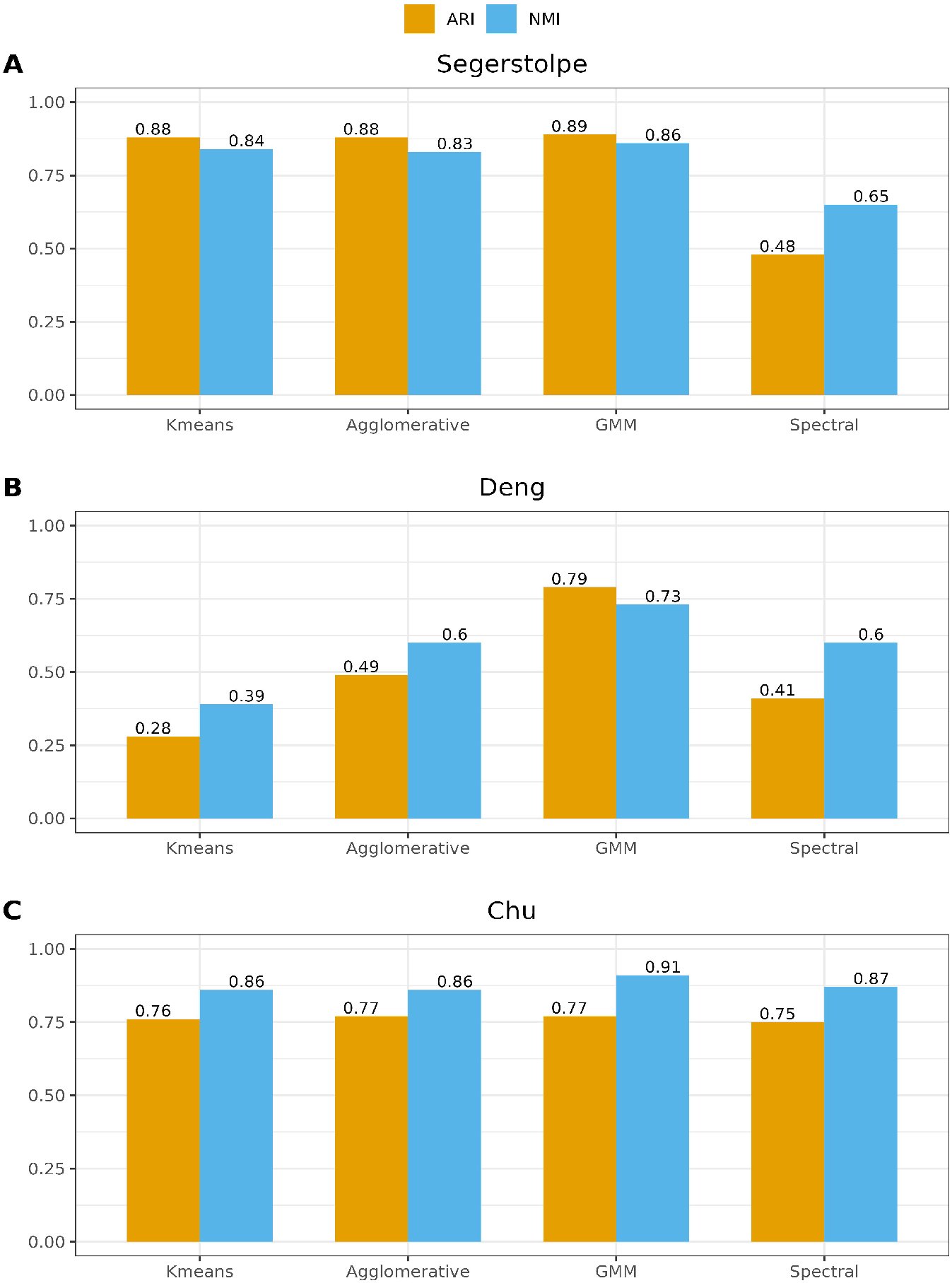
Shows the ARI and NMI scores of four different clustering algorithms using the feature space extracted by best performing dimensionality reduction method for each dataset.

## Discussion

Single cell sequencing technology has seen rapid development in recent years allowing to study similarities, or differences at the single cell level. In the last few years, a massive amount of scRNA-seq data has been uploaded to public databases such as the 10× Genomics, Human Cell Atlas and the Mouse Cell Atlas. One major challenge with the scRNA-seq data analysis is its high dimensionality and sparsity. The goal here is to extract maximum information from such big data by pruning redundant dimensions. Further, it is important to identify similarity and sparsity between different cells and finally assign cell types while minimizing computational cost. To this end, dimensionality reduction methods are widely used in scRNA-seq data analysis to tackle the issue of high dimensionality inherent in such datasets. For example, PCA is a linear method which uses independent linear combinations of original feature vectors and arranges them in descending order of their variance. ICA is another linear method, but instead of maximizing the variance, ICA minimizes dependencies among new features. ICA is well known for being able to identify independent sources which make up a signal. Advanced nonlinear approaches that preserve both global and local structures well in the lower dimension representation, such as t-SNE and UMAP, have recently been adopted for scRNA-seq data. Recent benchmarking study used VAE and DCA methods which are deep learning models based on the general autoencoder structure. The unsupervised clustering is performed to group similar cells and to find rare cell types as outliers. The performance of such clustering algorithms relies to a larger extend, on the outcome of the dimensionality reduction method.

We performed a benchmarking study of seven popular dimensionality reduction methods and four clustering algorithms widely used for scRNA-seq datasets. We used three different scRNA-seq datasets publicly available for this purpose. In addition, we compared our results with a similar study previously published by Xiang and colleagues [19]. Noteworthy, we noticed several differences in our observations when compared with Xiang’s paper. For example, our study revealed ICA method as the best performer for the *Segerstolpe* dataset. On the other hand, Xiang’s paper reported VAE method as the best performer for the same dataset. One possible explanation for this difference might be that Xiang and colleagues have always utilized only the first two dimensions of their reduced feature space for the sake of uniformity. In contrast, in our study we used varying number of reduced dimensions for clustering. Another possible reason for this difference might be that the dataset curators instructed to remove some 1305 cells from the dataset which were a result of noise, low quality, no cell, or multiple cells being captured in one droplet. These erroneous cells were not removed in the analysis reported by Xiang’s paper, albeit we excluded them in our analysis. Furthermore, probably for the same reasons, we observed higher ARI and NMI scores as compared to those previously reported in Xiang’s paper for all dimensionality reduction methods. Similarly, we found UMAP as the best performing dimensionality reduction method for the *Deng* dataset, while Xiang and colleagues reported PCA to be the best. Moreover, we noticed a huge increase in the ARI and NMI scores of the ICA method for this dataset. This may be explained by the fact that linear models such as ICA have neither any hyperparameters nor do they involve any stochastic mechanism. Lastly, for the Chu dataset, we observed a very drastic increment in accuracy of UMAP (i.e., ARI score increased from 0.15 to 0.502 and NMI score increased from 0.11 to 0.72). In addition, we noticed a decrease in accuracies for remaining dimensionality reduction methods for this dataset. Likely explanation for this difference might be that Xiang’s paper has utilized only 2 cell types (DEC and H1), while original dataset had seven cell types (DEC, H1, H9, EC, NPS, HFF, TB). We could not find any justification for excluding remaining 5 cell types in the Xiang’s study. We have not compared our analysis with that done by Tsuyuzaki and colleagues [20] as their main aim was to benchmark different variations of the PCA method, while the aim of our study was to overall compare dimensionality reduction methods from a wide range of categories and to elucidate whether certain types of categories perform consistently well or not.

Moreover, we benchmarked the performance of four clustering algorithms (k-means, agglomerative, GMM and spectral clustering) utilizing the latent space provided by the best performing dimensionality reduction method for each dataset. For instance, we used the features extracted by ICA method for the *Segerstolpe* dataset. Overall, we found no major differences in the performances of four clustering algorithms for this dataset. This indicates that ICA was able to appropriately minimizes dependencies among new features, which results in better cluster formation using any of the four algorithms. ICA is well known for being able to identify independent sources which make up a signal. Likewise, for the *Deng* and *Chu* datasets, we utilized the reduced dimensions extracted by UMAP and t-SNE methods, respectively. Both t-SNE and UMAP are advanced nonlinear approaches that preserve both global and local structures well in the lower dimension representation. Dimensionality reduction methods like t-SNE are designed to capture the data within 4 dimensions, while other methods used in our study do not have this restriction. Moreover, we iterated over a few dimensions of UMAP’s latent space to find the best dimensions and used them for benchmarking of the four clustering methods considered in our study. For the *Deng* dataset, the GMMs outperformed compared to other three clustering methods. This might be because GMMs are probabilistic and do soft assignment of clusters. They are even called the universal approximators of posterior distribution just as neural networks are called the universal approximators of functions. They can very accurately identify noncircular, anisotropicaly distributed and overlapping clusters. For *Chu* dataset, again we found no major differences in the performances of four clustering algorithms for this dataset. Though, GMMs marginally outperformed compared to other three clustering methods.

In summary, we conclude that the performance of different dimensionality reduction methods is dataset dependent. Noteworthy, t-SNE, UMAP and ICA performed well across the three datasets considered in the benchmarking study. The linear ICA method was the best performer for *Segerstolpe* dataset, while nonlinear methods UMAP and t-SNE best performed for *Deng* and *Chu* datasets, respectively. Moreover, the GMMs clustering algorithm performed consistently across different datasets. Contrary to our hypothesis, the deep learning methods VAE and DCA did not performed well in our study probably due to their sensitivity to hyperparameters and overfitting.

## List of Abbreviations

RPKM: Reads Per Kilobase of transcript per Million mapped reads
t-SNE: t-Student Stochastic Neighbor Embedding
UMAP: Uniform Manifold Approximation and Projection
GMM: Gaussian Mixture Model
PCA: Principal Component Analysis
VAE: Variational Autoencoder
DCA: Deep Count Autoencoder
ARI: Adjusted Random Index
NMI: Normalized Mutual Index
ZIFA: Zero Inflated Factor Analysis
ICA: Independent Component Analysis
EM: Expectation Maximization

## Acknowledgments

PY acknowledges the seed grant (project number I/SEED/PY/20200037) funded by the Indian Institute of Technology, Jodhpur.

## Author Contributions

VP performed the primary analysis; AKS contributed to figure preparation and manuscript writing; PY designed and supervised the study; all authors contributed to manuscript writing; all authors contributed by comments and approved the final manuscript.

